# The role of human superior colliculus in affective experiences during visual and somatosensory stimulation

**DOI:** 10.1101/2022.12.09.519812

**Authors:** Danlei Chen, Philip A. Kragel, Paul W. Savoca, Lawrence L. Wald, Marta Bianciardi, Tor D. Wager, Karen S. Quigley, Ajay B. Satpute, Lisa Feldman Barrett, Jordan E. Theriault

## Abstract

The superior colliculus is often studied for its role in visually guided behaviors, but research in non-human animals indicates it is a midbrain hub for processing sensory information from multiple domains, including interoception (which is associated with affect). We used ultra-high field 7-Tesla fMRI to extend this work to humans, modeling superior colliculus BOLD signal intensity during visual or somatosensory stimulation (N = 40 in each sensory modality), both under aversive and neutral affective intensity. As hypothesized, the superior colliculus showed increased BOLD signal intensity in the dorsal and ventral subregions during visual and somatosensory stimulation, respectively. The entire superior colliculus also showed increased BOLD signal intensity during aversive compared to neural conditions. The superior colliculus BOLD signal intensity also correlated with a preregistered set of brain regions involved in visual, somatosensory, and interoceptive processing.

## Introduction

The superior colliculus is the midbrain target of the optic nerve and has been studied extensively in terms of its functions in visual attention and visually guided behaviors (for review, see Krauzlis et al., 2013). For example, research using non-human vertebrates suggests that the superior colliculus plays an important role in visuomotor integration (for review, see Hagan et al., 2020; Wurtz & Albano, 1980) and in humans, the superior colliculus responds robustly to signals from the retina during visual stimulation (e.g., Schneider & Kastner, 2005), stimulus driven eye-movement (e.g., Krebs et al., 2010), memory-guided eye-movement and covert visual attention shift (e.g., Anderson & Rees, 2011), and visually guided body movement (e.g., Linzenbold & Himmelbach, 2012). Research with non-human vertebrates further suggests that the superior colliculus plays an important role in processing exteroceptive sensory signals beyond vision, including the auditory and somatosensory modalities, as well as multimodal integration in which sensory stimulation becomes stronger than the sum of each type of input (for review, see King, 2004). Studies of single-neuron recordings showed that neurons in the intermediate and deep layers of the superior colliculus respond to visual, auditory, and somatosensory stimulation, as well as the combinations of these inputs, whereas neurons in the superficial layers are organized into retinographic receptive fields and respond exclusively to visual stimulation (for review, see May, 2006). The primary goal of the present study is to examine the human superior colliculus during the processing of signals from sensory domains beyond visually related functions, which has yet to be explored in humans.

The multisensory capacity of the superior colliculus very likely includes regulating the body’s internal systems (called visceromotor control), in addition to the exteroceptive domains. Both structural and functional findings suggest that the superior colliculus plays a broader role in sensory processing that helps select appropriate skeletomotor actions to approach or avoid a stimulus (for review, see Gandhi & Katnani, 2011; Isa et al., 2021; Suzuki et al., 2019) as well as to regulate the associated visceromotor changes within the body that support those actions (e.g., Keay et al., 1988). For example, stimulating the superior colliculus in rodents produces orienting or defensive behaviors accompanied by large increases in blood pressure and heart rate, whereas lesions of the superior colliculus in rodents and primates diminish such responses in the face of potential threats (e.g., Blanchard et al., 1981; Keay et al., 1988; Maior et al., 2012; Redgrave & Dean, 1985; Sahibzada et al., 1986). Visceromotor activity that supports action is coupled with internal sense data from the body generated by that activity, called interoception (for discussion, see Barrett & Simmons, 2015; Craig, 2014; Kleckner et al., 2017; Quigley et al., 2021; Seth & Friston, 2016). This relationship led us to hypothesize that the superior colliculus may also process interoceptive sense data.

Consistent with this hypothesis, increases in visceromotor activity and the related changes in interoception are associated with more intense affective experience in humans (for review, see Critchley & Garfinkel, 2017), and correspondingly, brain imaging studies show increased BOLD signal intensity in the superior colliculus as human participants view images that evoke intense affective experience (e.g., Almeida et al., 2015; Kragel et al., 2021; Wang et al., 2020). A role for the superior colliculus in interoception (and by association, affective experience) is further supported by its anatomical connections with a range of cortical and subcortical structures known to be involved in interoception and visceromotor control (see Table 1). Specifically, the anterior cingulate cortex (e.g., Harting et al., 1991), the hypothalamus (e.g., Benevento & Fallon, 1975), and the periaqueductal gray (e.g., Beitz, 1989) are connected with the intermediate/deep layers of the superior colliculus, suggesting that these layers may be involved with sensing and regulating the internal systems of the body (interoception and visceromotor control, respectively), consistent with its established role in affect.

**Table 1.**
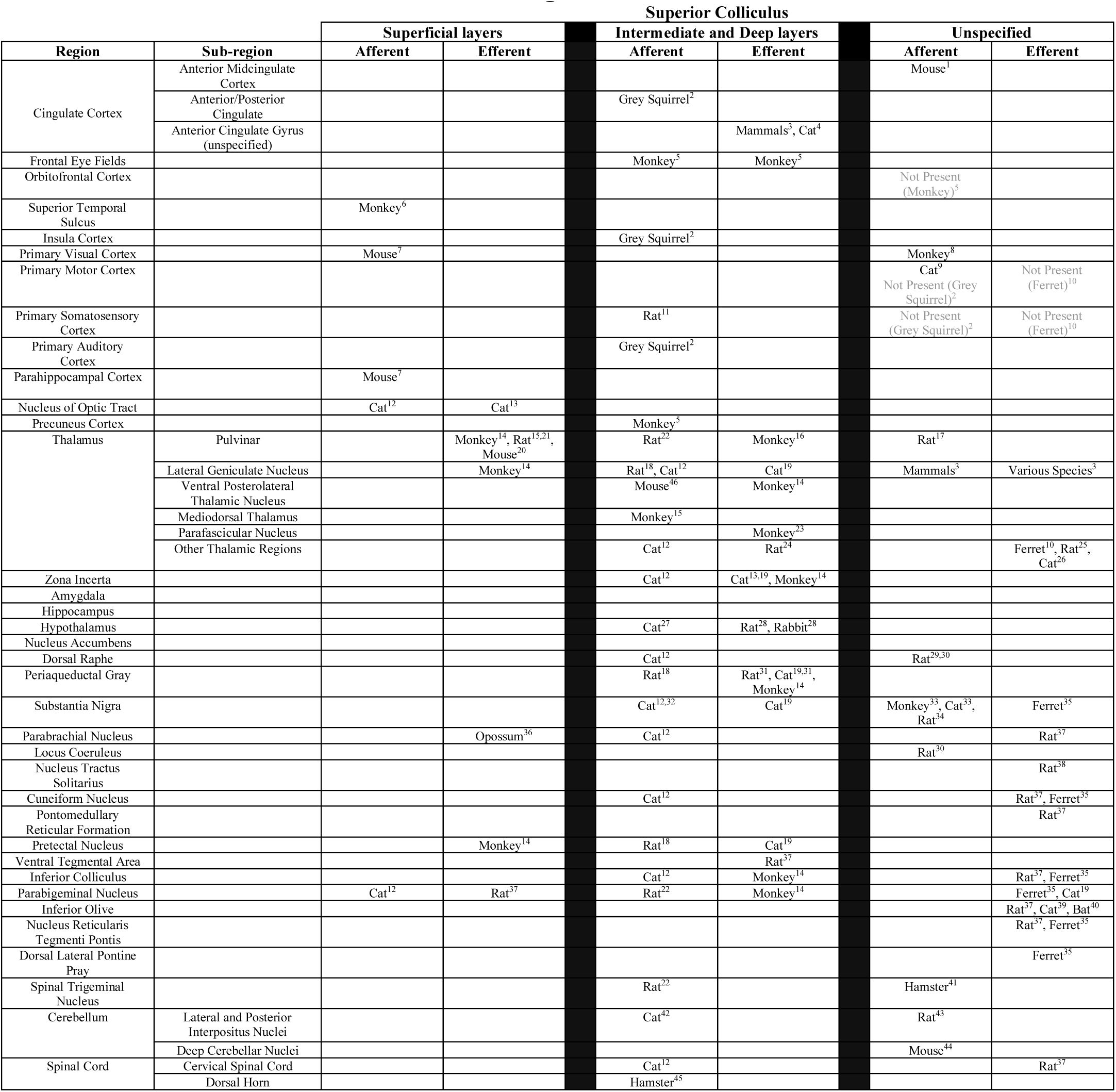
Monosynaptic afferent and efferent connections of the superior colliculus. Species in which connections were observed are listed in cells, with relevant citations listed below. Blank cells denote the *absence of evidence* for or against a connection, to the best of our knowledge. Cells labeled “not present” in gray text denote a study finding *evidence of absence*. In general, the superior colliculus projects to, and receives projections from, a wide range of cortical and subcortical regions important for exteroceptive (i.e., external sensory experience) and interoceptive (i.e., sensation from the viscera and internal milieu of the body; Craig, 2014) sensation. Of particular importance, the superficial and intermediate/deep layers of the superior colliculus show distinct patterns of anatomical connectivity. In the superficial layers, afferent and efferent connections primarily project to regions traditionally regarded as important for visual processing—e.g., afferent connections from the nucleus of optic tract and primary visual cortex and efferent connections to visual subregions of thalamus, including pulvinar and the lateral geniculate nucleus. By contrast, the intermediate and deep layers show broad connections related to both exteroception and interoception. The exteroceptively-relevant connections in the intermediate/deep layers include afferent connections from primary somatosensory cortex, primary auditory cortex, and pulvinar, as well as bidirectional connections to/from frontal eye field and the lateral geniculate nucleus. The interoceptively-relevant connections in the intermediate/deep layers include afferent connections from the insular cortex, mediodorsal thalamus, dorsal raphe, and parabrachial nucleus, as well as efferent connections to ventral tegmental area, and bidirectional connections to/from cingulate cortex, hypothalamus, periaqueductal gray, and substantia nigra. Moreover, the intermediate and deep layers of the superior colliculus also receive direct afferent inputs from the cerebellum and spinal cord. These observations confirm prior functional results, finding that the superficial layers of the superior colliculus are largely connected with vision-related brain regions, while the intermediate and deep layers of superior colliculus are connected with multimodal sensory regions, including multiple primary exteroceptive sensory regions and those within the interoceptive network (Kleckner et al., 2017). Superficial layers of SC include stratum zonale, stratum griseum superficiale, and stratum opticum. Intermediate layers include stratum griseum intermediate and stratum album intermediate. Deep layers include stratum griseum profundum and stratum album profundum (Baldwin et al., 2019). Reference legend: ^1^Fillinger et al., 2018; ^2^Baldwin et al., 2019; ^3^Harting et al., 1991; ^4^Sherman & Sprague, 1979; ^5^Leichnetz et al., 1981; ^6^Fries, 1984; ^7^Wang & Burkhalter, 2013; ^8^Collins et al., 2005; ^9^Harting et al., 1992; ^10^Manger et al., 2010; ^11^Hoffer et al., 2005; ^12^Edwards et al., 1979; ^13^Graham, 1977; ^14^Benevento & Fallon, 1975; ^15^Sommer & Wurtz, 2004; ^16^Benevento & Standage, 1983; ^17^Taylor et al., 1986; ^18^Beitz, 1989; ^19^Grofová et al., 1978; ^20^Gale & Murphy, 2018; ^21^Donnelly et al., 1983; ^22^Cadusseau & Roger, 1985; ^23^Harting et al., 1980; ^24^Linke et al., 1999; ^25^Krout et al., 2001; ^26^Coizet et al., 2007; ^27^Rieck et al., 1986; ^28^Fallon & Moore, 1979; ^29^Villar et al., 1988; ^30^Waterhouse et al., 1993; ^31^Ranagnano, 1993; ^32^Graybiel, 1978; ^33^Beckstead, 1983; ^34^Comoli et al., 2003; ^35^Doubell et al., 2000; ^36^Rafols & Matzke, 1970; ^37^Redgrave et al., 1987; ^38^van der Kooy et al., 1984; ^39^Saint-Cyr & Courville, 1982; ^40^Covey et al., 1987; ^41^Van Buskirk, 1983; ^42^Kawamura et al., 1982; ^43^Gayer & Faull, 1988; ^44^Benavidez et al., 2021; ^45^Rhoades, 1981; ^46^Doykos et al., 2020.

In this study, we examined the human superior colliculus during visual and somatosensory stimulation at two levels of affective intensity (aversive and neutral). In humans, the spatial limitations of non-invasive imaging methods, like fMRI, have made it difficult to measure superior colliculus BOLD signal intensity: the superior colliculus is about 6 mm in diameter, the size of a pea, while traditional 3-Tesla fMRI offers a voxel-wise resolution of 2-3mm. To solve this problem, we used ultra-high field 7-Tesla fMRI (1.1 mm isotropic) to examine changes in superior colliculus BOLD signal intensity and functional connectivity (for related methods, see Kragel et al., 2019, 2021; Satpute et al., 2013; Wang et al., 2020). During scanning, 80 human subjects completed five fMRI runs (120 trials in total), involving either visual stimulation or somatosensory stimulation of the left thumb (*sensory modality*; between subject; N = 40 in each sensory modality). Half of the trials for each subject were affectively aversive (i.e., negatively-valenced images or strong press on the left thumb) and half were affectively neutral (i.e., neutrally-valenced images or weak press on the left thumb; *affective intensity*; within-subject; see Methods; Fig S1).

We examined two hypotheses regarding superior colliculus BOLD signal intensity (from here, BOLD signal intensity), guided by previous anatomical and functional findings reviewed above. First, we hypothesized that dorsal and ventral *subregions* of the superior colliculus (corresponding to superficial and deep layers, respectively; see Methods) would interact with sensory modality, such that visual stimulation would elicit increased BOLD signal intensity in the dorsal subregion, and somatosensory stimulation would elicit increased BOLD signal intensity in the ventral subregion. Second, we hypothesized that the entire superior colliculus would show increased BOLD signal intensity during aversive (vs. neutral) affective stimulation in both sensory modalities.

We preregistered three more hypotheses regarding the superior colliculus’s functional connectivity with several distributed brain networks (for details, see https://osf.io/pa5b9/). First, we hypothesized that superior colliculus BOLD signal intensity during each modality of sensory stimulation would correlate with the BOLD signal intensity in the corresponding primary sensory network (i.e., the visual network during visual stimulation and the somatosensory network during somatosensory stimulation)^1^. Second, we hypothesized that the superior colliculus BOLD signal intensity would correlate with the visual network during somatosensory stimulation and with the somatosensory network during visual stimulation, based on the previous findings of multisensory processing in the superior colliculus from non-human animals (for review, see King, 2004). Finally, we hypothesized that the BOLD signal intensity in the superior colliculus would correlate with the BOLD signal intensity in the interoceptive network (Kleckner et al., 2017) during trials stimulating both visual and somatosensory modalities^2^. This final hypothesis was based on non-human animal research identifying monosynaptic anatomical connections between the superior colliculus and brain regions involved in interoception (see Table 1).

## Results

### Task-related changes in BOLD signal intensity in the superior colliculus

#### Dorsal superior colliculus showed greater increases in BOLD signal intensity during visual stimulation, and ventral superior colliculus showed greater increases in BOLD signal intensity during somatosensory stimulation

After preprocessing the BOLD signal intensity (see Methods) during stimulation in visual and somatosensory modalities, contrast maps were estimated in each participant for both aversive and neutral affective intensity. To ensure that the superior colliculus was properly aligned across all participants (even after normalization to standard MNI space), participant-specific superior colliculus masks were identified and warped to generate group-specific superior colliculus masks. Participant-level contrasts were then transformed to align based on group-specific superior colliculus masks (see Methods). Group-level whole-brain BOLD contrasts were modeled and visualized using these participant-level aligned contrasts (see Methods). Group-level whole-brain contrasts of the aligned superior colliculus showed expected patterns of BOLD signal intensity (e.g., visual cortex during visual stimulation; right primary somatosensory cortex during somatosensory stimulation of the left thumb; see Fig. S2). As hypothesized, the BOLD signal intensity increased in the superior colliculus relative to a baseline fixation in *both* visual and somatosensory stimulation and at both levels of affective intensity (Fig. 1).

**Fig 1.**
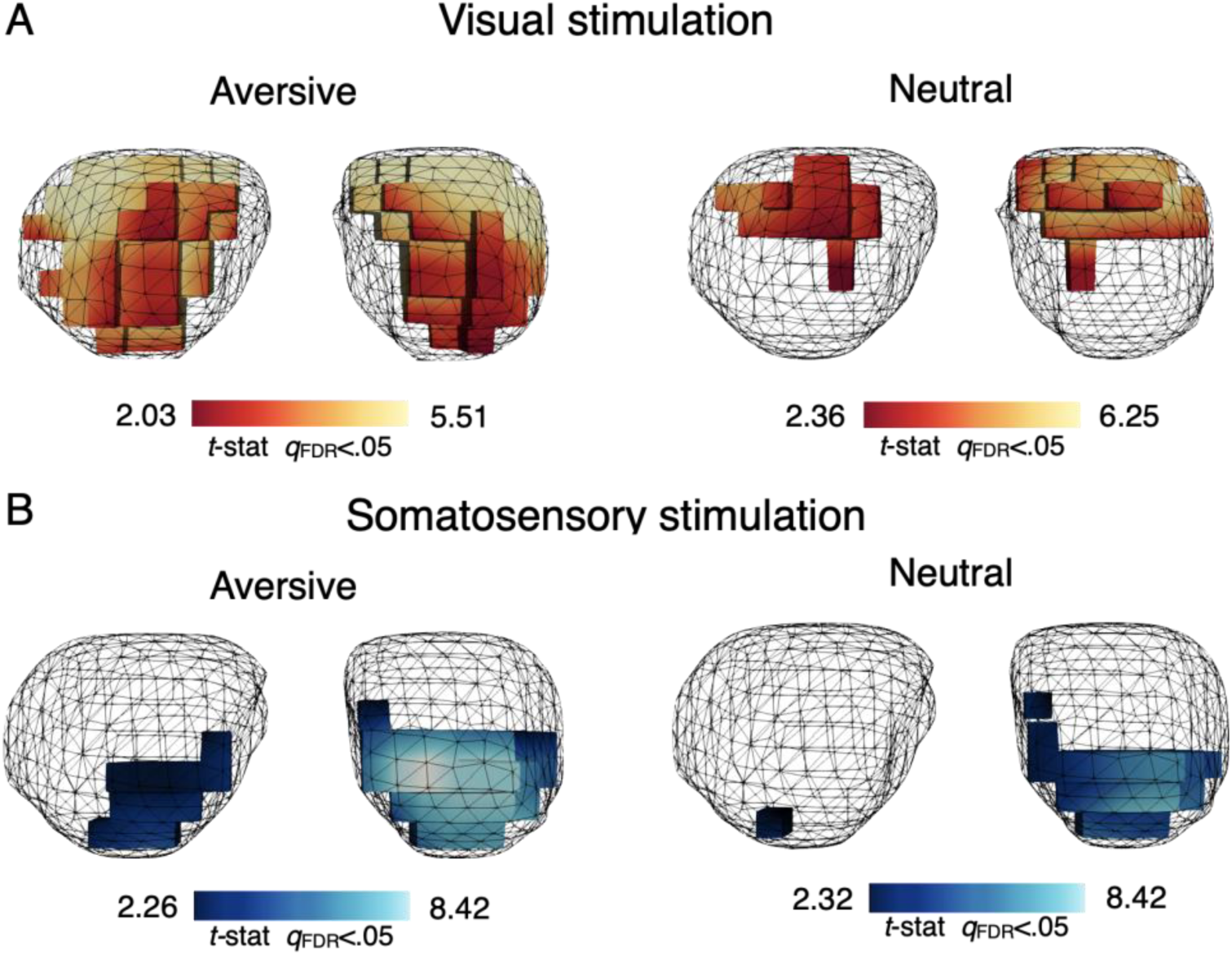
Group-level univariate analysis in the superior colliculus. Superior colliculus BOLD signal intensity for aversive and neutral affective intensity during visual (A; N = 40) and somatosensory (B; N = 40) stimulation. Activation refers to one-tailed t statistics (stimulation > baseline), thresholded at a voxel-wise cutoff of *q*_FDR_ < 0.05. Results are shown in 3-dimentional semi-transparent mesh of group-specific superior colliculus masks.

As hypothesized, visual and somatosensory stimulation was associated with distinct patterns of BOLD signal intensity within the superior colliculus. A multivoxel pattern analysis (MVPA) distinguished patterns of superior colliculus BOLD signal intensity associated with visual and somatosensory stimulation at above-chance accuracy (mean accuracy = 79.42% (chance accuracy = 50%), SD = 2.44%), using a linear support vector machine (SVM) classifier and leave-one-subject-out cross-validation (see Supplementary material for details). More specifically, a linear mixed effects model analysis (Fig. 2A) showed two important interactions: a significant interaction between subregion and sensory modality (Fig. 2B), *F_(1, 458.00)_* = 35.66, *P* < .001, and a significant interaction between laterality and sensory modality (Fig. 2C), *F_(1, 459.14)_* = 10.38, *P* < .001, such that visual stimulation was associated with greater BOLD signal intensity in bilateral dorsal superior colliculus (corresponding to the superficial layers), whereas somatosensory stimulation was associated with greater BOLD signal intensity in ventral, largely right-lateralized (contralateral to the left thumb, where pressure was applied) superior colliculus (corresponding to the intermediate and deeper layers).

**Fig 2.**
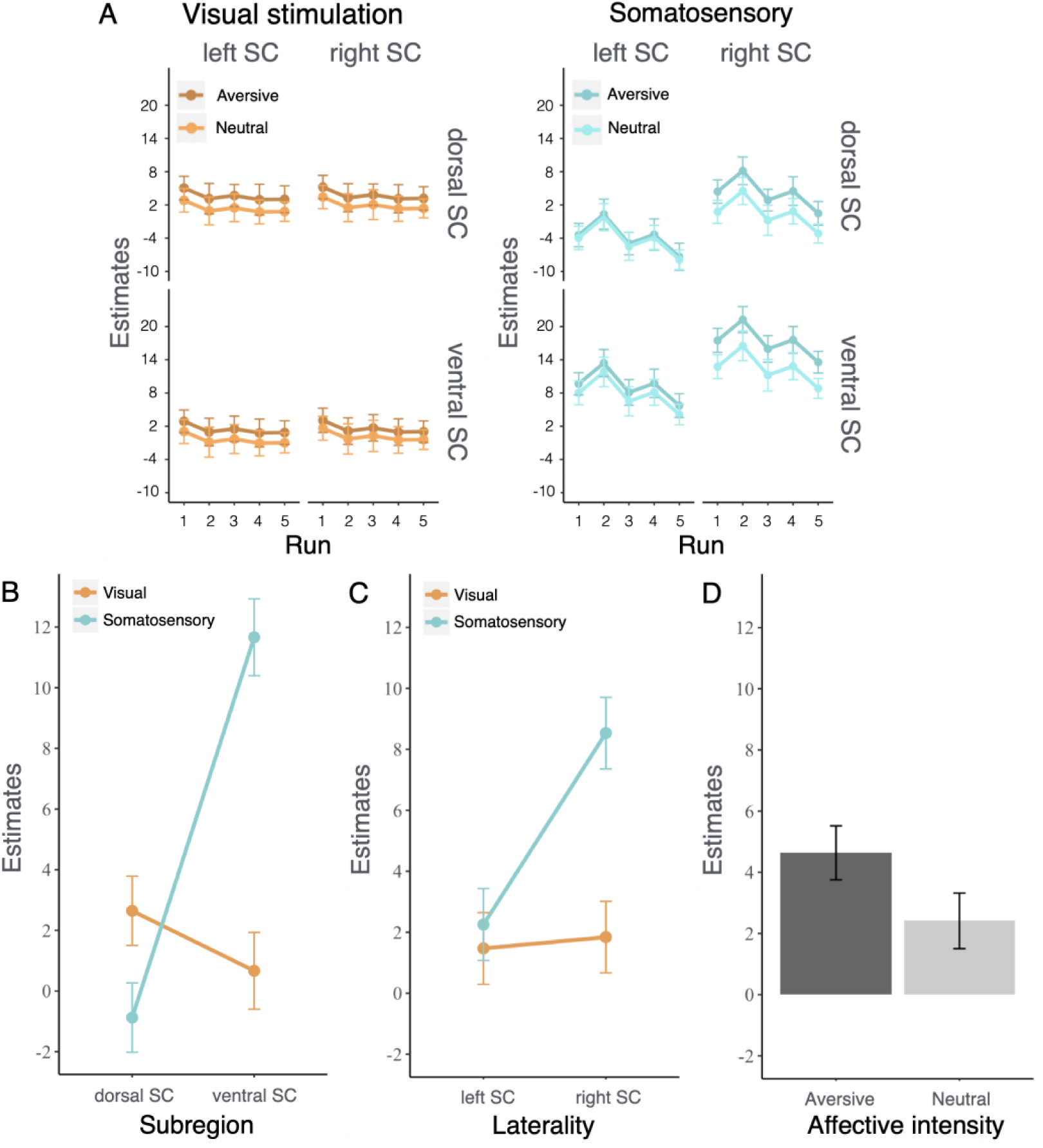
Group-level estimated marginal means from linear mixed effects model within the superior colliculus (see Methods for full model description). (A) Full comparisons of estimated marginal means between sensory modality (visual/somatosensory; N = 40 in each), run (1–5), subregion (dorsal/ventral), laterality (left/right), and affective intensity (aversive/neutral). (B) A significant main effect of subregion, *F_(1, 79.51)_* = 44.67, *P* < .001, and a significant interaction between subregion and sensory modality, *F_(1, 79.51)_* = 84.37, *P* < .001, such that visual stimulation elicited greater BOLD signal intensity in dorsal than ventral superior colliculus subregion and somatosensory stimulation elicited greater BOLD signal intensity in ventral than dorsal superior colliculus subregion. (C) A main effect of laterality, *F_(1, 159.68)_* = 23.02, *P* < .001, and a significant interaction between laterality and sensory modality, *F_(1, 159.68)_* = 18.17, *P* < .001, such that somatosensory stimulation elicited a greater difference in BOLD signal intensity between laterality (right > left) than visual stimulation. (D) A main effect of affective intensity, *F_(1, 66.79)_* = 5.40, *P* < .05, such that aversive affective stimulation elicited greater BOLD signal intensity than neutral affective stimulation. We also observed a main effect of sensory modality, such that total superior colliculus BOLD signal intensity was greater during somatosensory stimulation than during visual stimulation, *F_(1, **70.16)**_* = 6.25, ***P*** < .05. All other comparisons were non-significant. Error bars indicate ± l standard error. See Table Sl for complete model summary.

As an exploratory result, we also found that the ventral superior colliculus was involved in the processing of both visual and somatosensory stimulation with increased BOLD signal intensity in compared to baseline (visual > baseline: *t*_(39)_ = 2.13, *P* < .05; somatosensory > baseline: *t*_(39)_ = 6.38, *P* < .001). By contrast, dorsal superior colliculus was only involved in stimulation in the visual domain. That is, only visual stimulation was associated with increased BOLD signal intensity in dorsal superior colliculus compared to baseline (visual > baseline: *t*_(39)_ = 7.38, *P* < . 001), while somatosensory stimulation was not (somatosensory > baseline: *t*_(39)_ = -0.27, *P* > .05). These results are consistent with previous non-human vertebrate findings such that the deep layers of superior colliculus are multisensory while the superficial layers are only visual (see May, 2006).

#### Superior colliculus showed greater increases in BOLD signal intensity during aversive stimulation compared to affectively neutral stimulation

As hypothesized, aversive trials were associated with a greater increase in BOLD signal intensity throughout the superior colliculus, compared to neutral trials (Fig. 1). Results from the linear mixed effects model confirmed this observation with a main effect of affective intensity (Fig. 2D), *F_(1, 123.89)_* = 4.06, *P* < .05, such that the aversive affective stimulation was associated with greater BOLD signal intensity in the superior colliculus compared to neutral affective stimulation.

### Task-related changes in functional connectivity of the superior colliculus

#### The superior colliculus showed functional connectivity with structures within the corresponding sensory networks during visual and somatosensory stimulation

Whole-brain functional connectivity was estimated during both visual and somatosensory stimulation using concatenated trial-wise BOLD estimates within each subject (see Methods). The average estimate in the subject-specific superior colliculus mask (for the importance of subject-specific masks in functional connectivity, see Sohn et al., 2015) was correlated with all voxels in the brain across trials using a split-half bootstrapping procedure (1000 iterations) to assess the *reliability* of signal connectivity (see Methods). This technique was used to help prevent False Negative errors, which can be exacerbated by both stringent statistical thresholds (Yarkoni, 2009) and the noise inherent to imaging small structures, such as the superior colliculus (e.g., from partial volume effects), allowing us to set a more liberal statistical threshold to detect weak, but reliable connectivity. Reliability maps showed the percentage of iterations where superior colliculus BOLD signal intensity significantly correlated with each voxel at *P* < .05 in both split-half samples (Fig. 3S). In addition, significance thresholding was calibrated for each voxel by using a permutation-derived null distribution (Fig. 3; see Methods).

**Fig 3.**
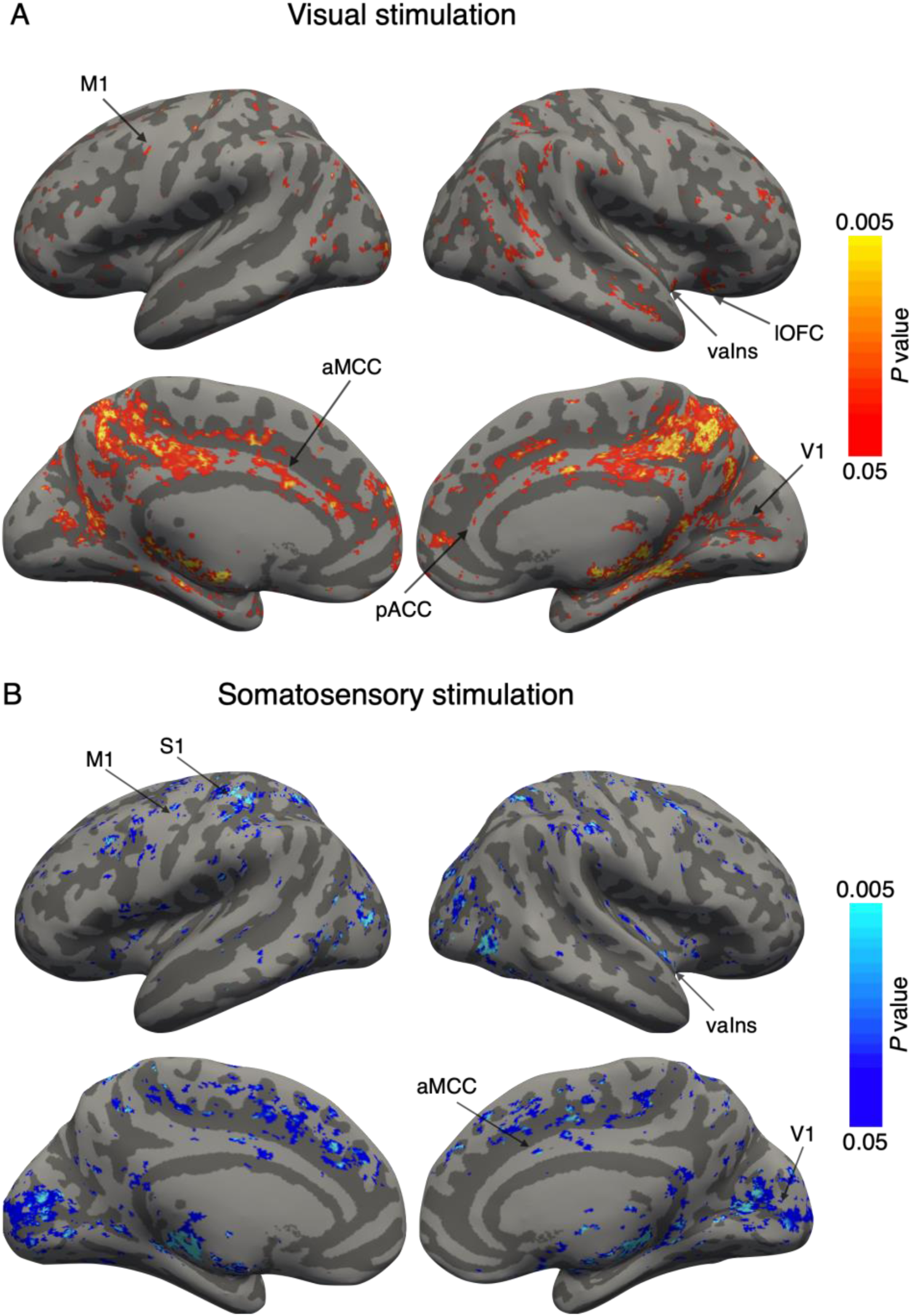
Group-level whole-brain reliability of functional connectivity with superior colliculus BOLD signal intensity during (A) visual (orange) and (B) somatosensory (blue) stimulation. Reliability was estimated based on the percentage of overlap in a bootstrapped split-half procedure (1000 iterations; see Methods). Thresholding for statistical significance was performed using a voxel-specific null-distribution, created by permutation (see Methods). Both panels report *P* values, scaled by -log_10_(*P*) and thresholded at *P* < .05 (equivalent of -log_10_(*P*) < 1.3). Cluster extent thresholding was set at k >= 20 voxels. Unthresholded 1-*P* value maps are available online: https://neurovault.org/collections/MKTVBWGR/. Percent coverage within ROIs of a whole-brain atlas (github.com/canlab) was calculated from the thresholded reliability maps and are available online (https://osf.io/pa5b9/). Abbreviations: AG: Angular Gyrus; lOFC: lateral orbitofrontal cortex; S1: primary somatosensory cortex; M1: primary motor Cortex; vaIns: ventral anterior insula; vmIns: ventral mid insula; aMCC: anterior mid cingulate cortex; pACC: pregenual anterior cingulate cortex; V1: primary visual cortex.

Consistent with our first preregistered hypothesis, during visual stimulation, superior colliculus BOLD signal intensity reliably correlated with the BOLD signal intensity in the visual network, including the primary visual cortex, lateral geniculate nucleus (LGN), and pulvinar (Fig. 3A & Fig. 4A). Likewise, during somatosensory stimulation, superior colliculus BOLD signal intensity reliably correlated with the BOLD signal intensity in the somatosensory network, including the primary somatosensory cortex and ventral posterolateral thalamic nucleus (VPL; Fig. 3B & Fig. 4B).

**Fig 4.**
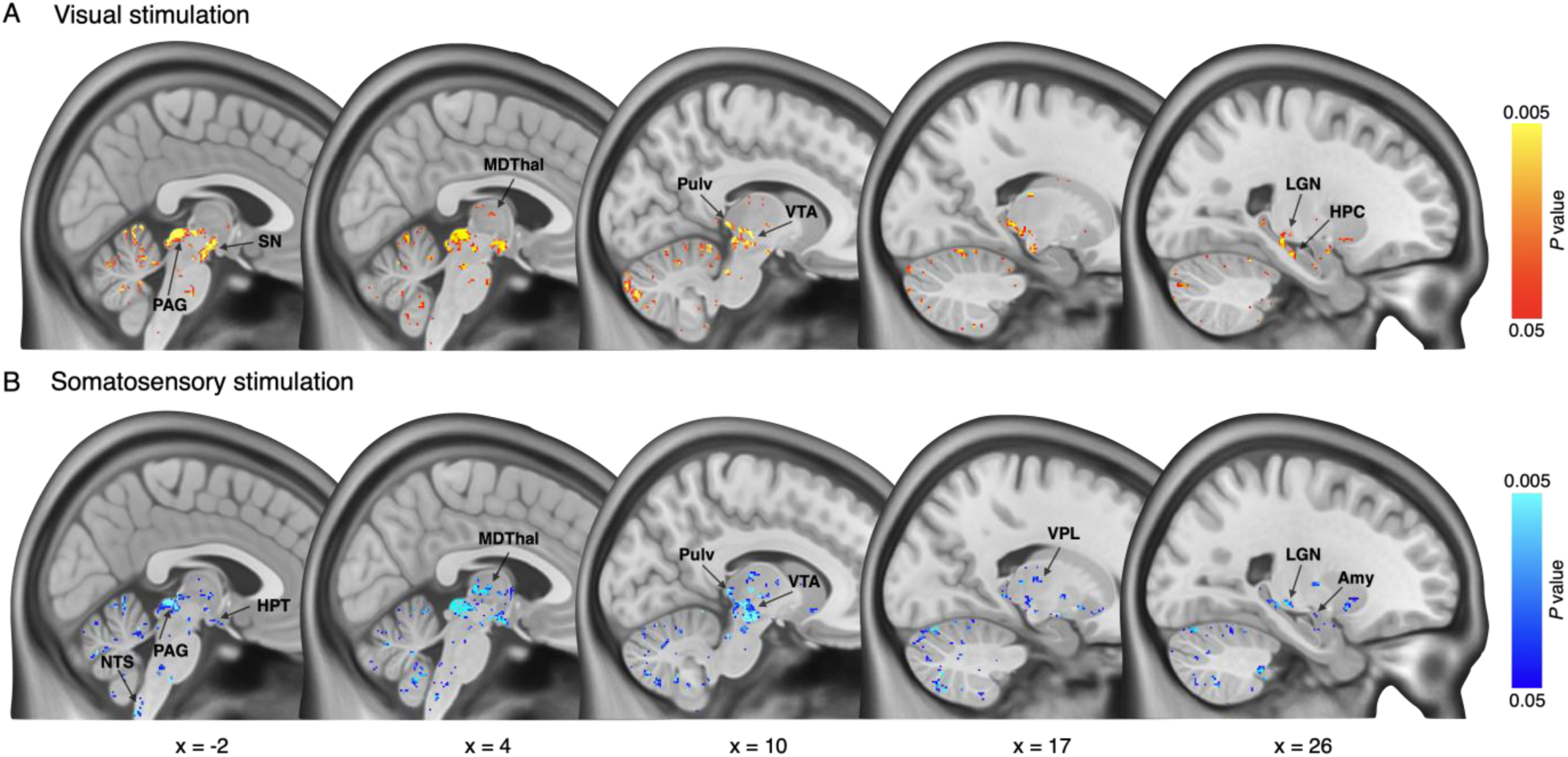
Group-level subcortical reliability of functional connectivity with superior colliculus BOLD signal intensity during (A) visual (orange) and (B) somatosensory (blue) stimulation. Reliability was estimated based on the percentage of overlap in a bootstrapped split-half procedure (1000 iterations; see Methods). Thresholding for statistical significance was performed using a voxel-specific null-distribution, created by permutation (see Methods). Both panels report *P* values, scaled by -log_10_(*P*) and thresholded at *P* < .05 (equivalent of -log_10_(*P*) < 1.3). Cluster extent thresholding was set at k >= 20 voxels. Abbreviations: Amy: amygdala; HPC: hippocampus; HPT: hypothalamus; LGN: lateral geniculate nucleus; MDThal: mediodorsal thalamus; NTS: nucleus of solitary tract; SN: substantia nigra; Pulv: pulvinar; PAG: periaqueductal gray; VTA: ventral tegmental area; VPL: ventral posterolateral thalamic nucleus.

#### The superior colliculus showed some evidence of cross-modality functional connectivity during visual and somatosensory stimulation

Consistent with our second preregistered hypothesis, we observed some evidence of cross-modality connectivity during visual and somatosensory stimulation. During somatosensory stimulation, cross-modal connectivity was clearly observed, where superior colliculus BOLD signal intensity reliably correlated with the BOLD signal intensity in visual network regions, including the primary visual cortex, LGN, and pulvinar (Fig. 3A & 4A). During visual stimulation, cross-modal connectivity was relatively weaker, where superior colliculus BOLD signal intensity reliably correlated with the BOLD signal intensity in a somatosensory network region, the VPL, but not in the primary somatosensory cortex at conventional levels of thresholding (Fig. 3B & 4B), although some connectivity with the primary somatosensory cortex was observed with a more liberal threshold (see Fig. S3).

#### Superior colliculus showed reliable functional connectivity with the interoceptive and allostatic network during both visual and somatosensory stimulation

Consistent with our final preregistered hypothesis, superior colliculus BOLD signal intensity correlated with the BOLD signal intensity in several key cortical and subcortical regions in the interoceptive network (e.g., Berntson & Khalsa, 2021; Evrard, 2019; Gianaros & Sheu, 2009; Harper et al., 2003; Sepulcre et al., 2012; Wager et al., 2009; Zunhammer et al., 2021; see, Kleckner et al., 2017). During visual stimulation, reliable cortical connectivity with the superior colliculus included primary visceromotor regions (e.g., anterior mid cingulate cortex (aMCC) and pregenual anterior cingulate cortex (pgACC)), secondary visceromotor region (e.g., primary motor cortex, which contains visceromotor maps; see Levinthal & Strick, 2012), and multisensory regions (e.g., ventral anterior insula and lateral orbitofrontal cortex; Fig. 3A). During visual stimulation, reliable subcortical connectivity with the superior colliculus included the amygdala (including the central nucleus), hippocampus (throughout the long axis; no septal nuclei), thalamus (including the mediodorsal nucleus), hypothalamus (including ventromedial hypothalamus), substantia nigra (SN), ventral tegmental area (VTA), the entire PAG, and the cerebellum (including area V, I_IV, Vermis IX, and Crus II; Fig. 4A & Fig. 5).

**Fig 5.**
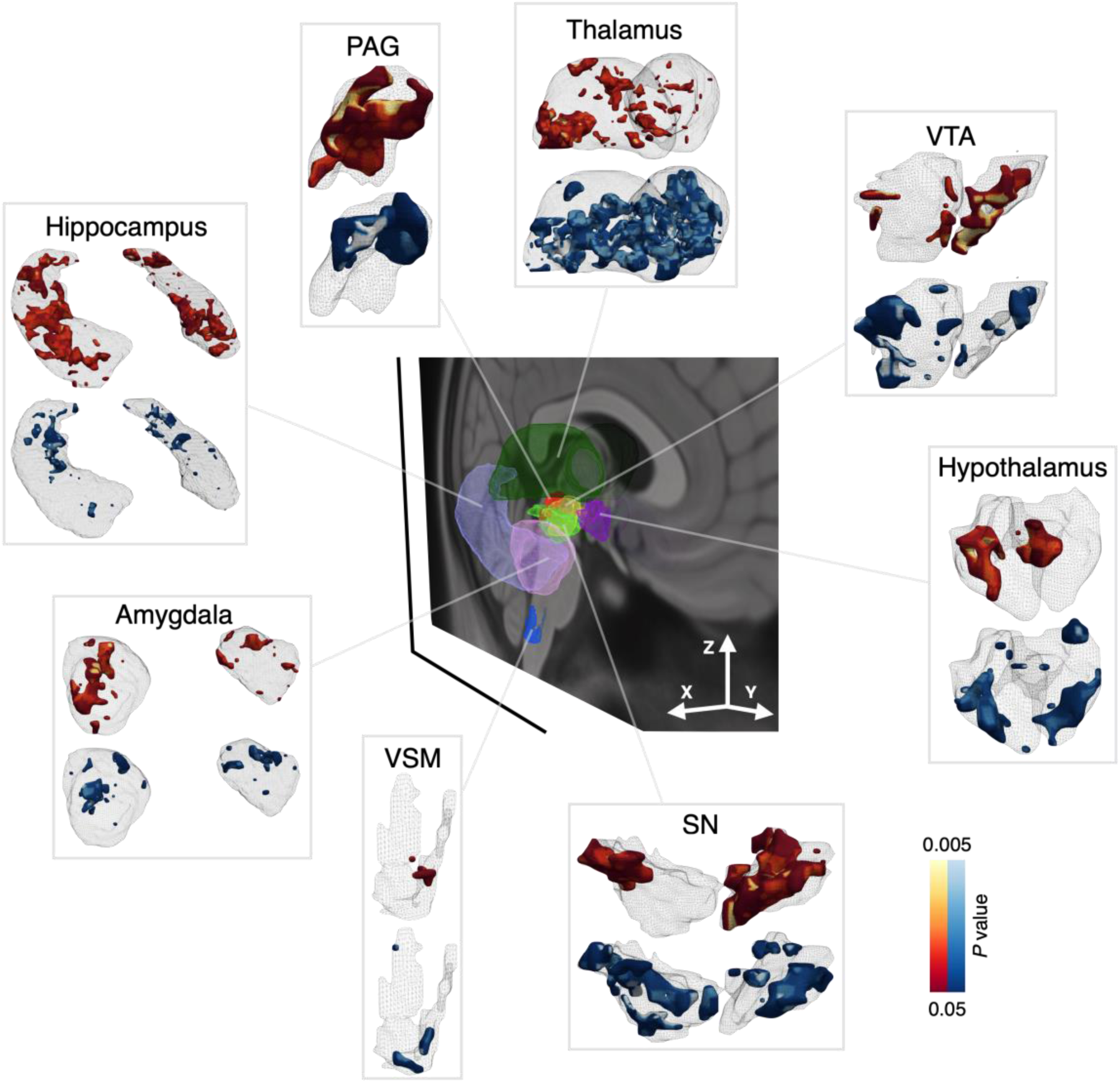
3D visualization of reliability of group-level functional connectivity in selected subcortical regions during visual (orange) and somatosensory (blue) stimulation. The middle panel depicts ROI location and orientation, overlaid onto a midline (x = 0) sagittal brain slice (slightly tilted, as indicated by the x/y/z coordinates), with 3D semi-transparent ROI meshes shown for the right hemisphere only. The radiated panels depict semi-transparent 3D meshes for each ROI, in the same orientation as in panel A, and enlarged for illustrative purposes (ROIs are shown at the appropriate relative sizes in the middle panel). ROIs display the same results as displayed in Fig. 4. Reliability was estimated based on the percentage of overlap in a bootstrapped split-half procedure (1000 iterations; see Methods). Thresholding for statistical significance was performed using a voxel-specific null-distribution, created by permutation (see Methods). All panels report *P* values, scaled by -log_10_(*P*) and thresholded at *P* < .05 (equivalent of -log_10_(*P*) < 1.3). Abbreviations: PAG: periaqueductal grey; SN: substantia nigra; VTA: ventral tegmental area; VSM: viscero-sensory-motor complex.

During somatosensory stimulation, reliable cortical connectivity with superior colliculus was similar to that observed during visual stimulation (i.e., anterior mid cingulate cortex, primary motor cortex, and ventral anterior insula; Fig. 3A), except for the lack of connectivity with pregenual anterior cingulate cortex and lateral orbitofrontal cortex. Subcortical connectivity was also observed in some of the same subcortical regions as that during visual stimulation (i.e., amygdala, hippocampus, mediodorsal thalamus, hypothalamus, SN, VTA, PAG, and cerebellum areas V, I_IV). Some differences were observed in the following subcortical regions – PAG connectivity was limited to the dorsal subregion; hippocampus connectivity was mostly in the tail; and connectivity to additional cerebellum areas VIIIA and Crus I (Fig. 4B & Fig. 5).

## Discussion

The present research used high-resolution 7-Tesla fMRI to examine the function of human superior colliculus. We observed a clear distinction between the dorsal and ventral subregions in the superior colliculus during visual and somatosensory stimulation in humans for the first time to our knowledge. This result aligns with the anatomical and functional findings in non-human vertebrates, in which the superior colliculus was shown to play a role in somatosensory processing in addition to its well-established role in vision-related processing (May, 2006). Previously, 3-Tesla fMRI studies in humans observed a similar distinction between the dorsal and ventral subregions of the superior colliculus in visually guided behaviors, where a relatively dorsal subregion was associated with oculomotor behaviors (i.e., saccades) and a relatively ventral subregion was associated with visually-guided somatomotor behaviors (i.e., arm reach; Himmelbach et al., 2013; Linzenbold & Himmelbach, 2012). The low resolution used in this research may have hindered a clear separation of superior colliculus from surrounding midbrain structures (e.g., PAG) and any BOLD signal from the cerebrospinal fluid (see Sclocco et al., 2018). By contrast, the present research more precisely resolved superior colliculus subregions based on its layered structure and provided results that are consistent with non-human animal research and our own preregistered hypotheses.

In addition to showing that the human superior colliculus is involved in processing exteroceptive sensory information, we also showed that BOLD signal intensity in the superior colliculus was modulated by affective intensity in both visual and somatosensory stimulation, consistent with prior human findings in the visual domain (e.g., Wang et al., 2020a) as well as prior non-human vertebrate findings about the importance of the superior colliculus in approach and avoidance behavior and visceromotor regulation (e.g., Keay et al., 1988; Sahibzada et al., 1986). These results are consistent with the hypothesis that the multisensory signal processing within the superior colliculus might extend to interoceptive signals, as affective experience is linked to the changes in interoceptive sensation (Critchley & Harrison, 2013). This hypothesis is further supported by the observed functional connectivity between the superior colliculus and several cortical and subcortical structures within the interoceptive network (e.g., aMCC, pgACC, and PAG; see Kleckner et al., 2017). Particularly, the PAG is known to be important for autonomic control of visceromotor activity (for review, see Silva & McNaughton, 2019), and it is directly bordered by the deep layers of superior colliculus. It will be important for future research to investigate the relationship between the PAG and deep layers of superior colliculus during interoceptive processing in humans (for comparison between PAG and deep superior colliculus functions in non-human animals, see Bittencourt et al., 2005; Dampney et al., 2013; de Almeida et al., 2006).

The results of task-based functional connectivity we observed were also generally consistent with the non-human vertebrate tract-tracing findings used to guide our hypotheses (Table 1). In addition, we identified novel functional connectivity between the superior colliculus and some structures, such as SN (not observed in prior resting state connectivity analyses, see Cauzzo et al., 2022; but consistent with anatomical evidence in non-human vertebrates, see Grofová et al., 1978). Some reliable patterns of functional connectivity also diverged from the tract-tracing findings in non-human vertebrates, however. For example, monosynaptic anatomical connections between the superior colliculus and the hippocampus and between the superior colliculus and the amygdala have not been observed (see Foreman & Stevens, 1987; Linke et al., 1999) yet we observed reliable functional connectivity between these regions. Functional connectivity between the superior colliculus and the hippocampus may reflect a second order connection, such as through the hypothalamus, which has connections to both regions (see Insausti et al., 1987; Veazey et al., 1982). Functional connectivity between the superior colliculus and the amygdala might reflect an ascending connection from superior colliculus through the pulvinar, which has been documented to project to the amygdala in both humans and non-human animals (see Benevento & Fallon, 1975; Huang et al., 2022; Koller et al., 2019). In addition, there is also the possibility that hippocampal and amygdala connections with superior colliculus might also be novel in humans, suggesting opportunities for further discovery.

Together, our results showed that the superior colliculus is involved in the processing of multiple sensory modalities, including vision, somatosensation, and possibly interoception. Consistent with this idea, the superior colliculus was found to process additional species-specific sensory modalities, such as the signals involved in infrared vision (rattlesnakes; Hartline et al., 1978), electroception (fish; Bastian, 1982), magnetoreception (rats; Němec et al., 2001), and echolocation (bats; Valentine & Moss, 1997) (for review, see Isa et al., 2021). When combined with these findings, our results suggest that the superior colliculus may be a vital multimodal integration hub for multiple channels of sensory information, considerably broadening its computational role in humans beyond the empirical focus on its vision-related functions. Indeed, most brain regions that were originally assumed to have a relatively modular function have now been shown to be involved in more diverse functions, including the primary visual cortex (e.g., Liang et al., 2013) and the primary motor cortex (e.g., Graziano, 2016; Levinthal & Strick, 2012). Future research might therefore explore the role of the human superior colliculus in supporting more complex tasks that require coordination and integration across several sensory and motor modalities, aligned with an emerging perspective in non-human vertebrate research (see Basso et al., 2021; Krauzlis et al., 2013; Merker, 2007).

## Methods

### Subjects

140 subjects were recruited, of which 41 withdrew participation or ended the scan session before starting the present task, leaving a total N of 99. Of this sample, 50 subjects completed the task in the visual modality, and 49 completed the task in the somatosensory modality. After data quality exclusions (detailed below), the sample in both visual and somatosensory modalities consisted of 40 subjects, with 167 runs in total for each task (average number of runs included per subject = 4.175).

Individual runs were excluded on the basis of poor quality (assessed by visual inspection) or high motion. In the visual modality, 43 runs (across 12 subjects) were excluded for poor image quality (issues included, for example, registration-failures caused by subject motion), and 33 runs (across 12 subjects) were removed for high motion (> .5mm framewise displacement in > 20% of run TRs). In the somatosensory modality, 25 runs (across 6 subjects) were excluded for poor image quality, and 35 runs were removed (across 12 subjects) for high motion. Given these exclusion criteria, 10 subjects entirely removed from the visual modality, and 9 subjects were entirely removed from the somatosensory modality.

The final sample of 80 subjects (M_age_ = 26.58 years, SD = 5.87 years, 33 female, 47 male) were recruited from the greater Boston area. All subjects were between 18 and 40 years old, were right-handed, had normal or corrected to normal vision, were not pregnant, were fluent English speakers, and had no known neurological or psychiatric illnesses. Subjects were excluded from participating in the study if they were claustrophobic or had any metal implants that could cause harm during scanning. All subjects provided written informed consent and study procedures were completed as approved by the Partners’ Healthcare Institutional Review Board.

### Experimental Paradigm

In a probabilistic avoidance learning task (see Fig. S1), subjects associated visual cues with aversive or neutral stimuli. The task consisted of 5 runs, with 24 trials in each. Only the visual or somatosensory stimulation period (3 secs. for visual task and 1 sec for somatosensory; see Univariate Analysis below) was relevant to the main research question of this paper. The avoidance learning component of the task was irrelevant but is reported in full below to provide context.

Each trial started with the presentation of two visual cues: a circle and a triangle. For 50% of trials, subjects chose a shape, and in the remaining 50% of trials a shape was randomly chosen for them. This design ensured that subjects were exposed to the outcome of both shapes. Subjects were instructed to avoid choosing the shape that led to aversive outcomes and that the level of aversiveness associated with each shape could change over time. After the choice, either neutral or aversive reinforcement was delivered. In the visual modality, an IAPS image (Lang et al., 1997), normed to be neutral or aversive valence, was presented on the screen. In the somatosensory modality, mechanical pressure was delivered to the bed of left thumb at medium (i.e., neutral) or high (i.e., aversive) pressure. Reinforcement rates for each shape were pseudorandomly chosen from 4 predetermined random walks between 20% and 80%, with outcomes determined randomly on every trial.

For visual reinforcement, the same pre-selected set of IAPS images were used for all subjects with valence rated on a scale of 1–10. Neutral images had an average normative valence of 5.52 (SD = .57) and arousal of 3.16 (SD = .57), and aversive images had an average normative valence of 2.93 (SD = .67) and arousal of 4.75 (SD = .58). Aversive and neutral images are different significantly in both valence (*t*_(42.85)_ = 9.30, *P* < .001) and arousal (*t*_(40.86)_ = -13.72, *P* < .001). For somatosensory reinforcement, neutral mechanical pressure was approximately 3 kg/cm^2^, and aversive mechanical pressure was approximately 5 kg/cm^2^. The pressure device was adjusted to each subject to ensure that the 2 levels were distinguishable and that neither was severely uncomfortable.

### fMRI Data Acquisition and Preprocessing

Gradient-echo echo-planar imaging BOLD-fMRI was performed on a 7-Tesla Siemens MRI scanner. Functional images were acquired using GRAPPA-EPI sequence: echo time = 28 ms, repetition time = 2.34 s, flip angle = 75°, slice orientation = transversal (axial), anterior to posterior phase encoding, voxel size = 1.1 mm isotropic, gap between slices = 0 mm, number of slices = 123, field of view = 205 × 205 mm^2^, GRAPPA acceleration factor = 3; echo spacing = 0.82 ms, bandwidth = 1414 Hz per pixel, partial Fourier in the phase encode direction = 7/8. A custom-built 32-channel radiofrequency coil head array was used for reception. Radiofrequency transmission was provided by a detunable band-pass birdcage coil.

Structural images were acquired using a T1-weighted EPI sequence, selected so that functional and structural images had similar spatial distortions, which facilitated co-registration and subsequent normalization of data to MNI space. Structural scan parameters were: echo time = 22 ms, repetition time = 8.52 s, flip angle = 90°, number of slices = 126, slice orientation = transversal (axial), voxel size = 1.1 mm isotropic, gap between slices = 0 mm, field of view = 205 × 205 mm^2^, GRAPPA acceleration factor = 3; echo spacing = 0.82 ms, bandwidth =1414 Hz per pixel, partial Fourier in the phase encode direction = 6/8. fMRI data were preprocessed using FMRIPREP (Esteban et al., 2019), a Nipype (Gorgolewski et al., 2011) based tool. Spatial normalization to the ICBM 152 Nonlinear Asymmetrical template version 2009c was performed through nonlinear registration with the antsRegistration tool of ANTs v2.1.0 (Avants et al., 2008; Gorgolewski et al., 2011), using brain-extracted versions of both T1w volume and template. Brain tissue segmentation of cerebrospinal fluid (CSF), white-matter (WM) and gray-matter (GM) was performed on the brain-extracted T1w using FAST (FMRIB’s Automated Segmentation Tool; FSL v5.0.9; Y. Zhang et al., 2001). Functional data was slice time corrected using 3dTshift from AFNI v16.2.07 (Cox, 1996) and motion corrected using MCFLIRT (FSL v5.0.9; Jenkinson et al., 2002). This was followed by co-registration to the corresponding T1w using boundary-based registration (Greve & Fischl, 2009) with six degrees of freedom, using FLIRT (FMRIB’s Linear Image Registration Tool; FSL v5.0.9; Jenkinson et al., 2002; Jenkinson & Smith, 2001). Motion correcting transformations, BOLD-to-T1w transformation and T1w-to-template (MNI) warp were concatenated and applied in a single step using ants ApplyTransforms (ANTs v2.1.0) using Lanczos interpolation.

### Superior Colliculus Localization and Alignment

To account for individual variability in superior colliculus alignment to MNI space, we adapted a method from previous studies that localized the periaqueductal gray (PAG; Kragel et al., 2019; Satpute et al., 2013) and automated a procedure to locate superior colliculus in each subject. This procedure began with a hand-drawn superior colliculus mask template which was subsequently masked to include voxels in a subject-specific grey and white matter probability masks (> 25%) and exclude voxels in a subject-specific CSF probability mask (> 50%). This mask was further refined to exclude voxels in a subject-specific mask of the PAG and cerebral aqueduct (for procedure, see Kragel et al., 2019), large model residuals from the univariate subject-level analysis (> 85%; see Univariate Analysis below), and any lingering small clusters after the application of other exclusion criteria (< 10 voxels). Subject-specific superior colliculus masks were transformed to align all subject whole-brain data in a common space, correcting any misalignment that occurred in the transformation to MNI space. For visual and somatosensory groups separately, subject-specific superior colliculus masks were aligned using DARTEL (Ashburner, 2007), and the group-level transformations used in this alignment were then applied to subject-specific whole-brain contrast images.

### Univariate Analysis

To estimate BOLD signal intensity during visual and somatosensory modalities, a general linear model was applied to each subject’s preprocessed functional time series (GLM; FSL v5.0.9). First level models were estimated for each run in each subject. For the visual modality, the entire visual presentation duration (3 secs.) was modeled as a regressor. For the somatosensory modality, the onset of pressure delivery time (1 sec.) and the rest of pressure stimulation time (2 secs.) were modeled separately, consistent with prior studies of pain stimulation using similar experimental design (Roy et al., 2014). In both visual and somatosensory modalities, aversive and neutral trials were modeled separately. All regressors were convolved with a double gamma hemodynamic function. A visual cue and associated decision-making period preceded visual/somatosensory stimulation (5.5 secs. jittered) and was modeled, as well (but is not discussed). In all runs of both modalities, regressors were modeled in relation to the implicit baseline of the fixation cross. Nuisance regressors included a run intercept, motion (i.e. translation/rotation in x/y/z planes, and their mean-centered squares, derivatives, and squared derivatives), aCompCor components (5 CSF, 5 WM; Muschelli et al., 2014), a discrete cosine transformation set with a minimum period of 120 seconds, and spike regressors (> 0.5mm framewise displacement; Satterthwaite et al., 2013). Due to a programming error that omitted some timing information for certain trials when no choice responses were made, TRs for a small subset of trials were removed with spike regressors (0.6% of all trials). The entire run was dropped from analysis if > 20% of its TRs were excluded via spike regressors.

As discussed in Superior Colliculus Localization and Alignment above, first-level whole-brain contrasts were warped to align superior colliculus across subjects, then smoothed using a 3 mm FWHM Gaussian kernel. First-level contrasts were averaged across runs within each subject to create run-level whole-brain contrasts, which were in turn used to estimate group-level contrasts via ordinary least square (OLS) estimation using one-sample t*-*tests. Group-level whole-brain contrast results were thresholded by false discovery rate (*q*_FDR_ < 0.05), with a minimum cluster extent 20 voxels. Group-level voxel-wise superior colliculus results were also thresholded by false discovery rate (*q*_FDR_ < 0.05), but no minimum voxel extent was applied.

Additional voxel-wise analyses examined superior colliculus subregions. Non-human animal research has found sensory subregions in superficial to deep layers of the superior colliculus (e.g., Chalupa & Rhoades, 1977); however, we did not make direct distinctions between the anatomical layers of the superior colliculus because (a) superior colliculus layers are very thin, and difficult to delineate, even at 1.1mm resolution, and (b) the scanning direction of fMRI acquisition on the anterior-posterior plane (AP) could introduce partial volume effects, which would further reduce our ability to detect differences in thin superior colliculus layers in the dorsal-ventral and lateral-medial directions. Instead, we separated the superior colliculus into dorsal and ventral subregions by dividing the ROI in half. We hypothesized that the dorsal half (corresponding to superficial superior colliculus layers) would show increased BOLD signal intensity in the visual modality, and that the ventral half (corresponding to the deep layers) would show increased BOLD signal intensity in the somatosensory modality. This analysis approach unfortunately omits an investigation of the superior colliculus intermediate layers (which are well-connected with the hypothalamus, mediodorsal thalamus, and anterior cingulate cortex, among other regions; see Table 1); however, as noted above, our data is likely not to have high enough resolution to distinguish and compare specific superior colliculus layers.

Superior colliculus ROI analyses used a mixed effects linear model to model the fixed effects of run (1-5), laterality (left/right), subregion (defined as the upper and lower halves of superior colliculus divided by the midline; see above), affective intensity (aversive/neutral), interactions between affective intensity and laterality and between affective intensity and subregion, and interactions between all the mentioned fixed effects and sensory modality (visual/somatosensory). The remaining fixed effects and interactions were uninterpretable or not of theoretical interest and were omitted from the model. By-subject random effects included random intercepts and random slopes for all fixed effects listed above, except for the interactions with sensory modality, which was a between-subjects variable. In each subject-specific superior colliculus mask, voxel-wise estimates from unsmoothed first-level contrasts were averaged within run (maximum 5 per subject), laterality (2 per run), subregion (2 per run), and aversiveness (2 per run), providing a maximum of 20 estimates per subject. These averages were entered into a linear mixed effects model in R (R Core Team, 2016), using the *lmer4* package (Bates et al., 2015), and degrees of freedom were approximated by the Scatterhwaite method, as implemented in the *lmerTest* package (Kuznetsova et al., 2017). Marginal condition means were estimated using the *emmeans* package (Lenth, 2019). The code was entered as follows:

model <- lmer(mean_signal ∼ (run + laterality + subregion + affective_intensity + laterality : affective_intensity + subregion : affective_intensity)*sensory_modality+ (run + laterality + subregion + affective_intensity + laterality : affective_intensity + subregion : affective_intensity | subject), data=df)
marginal_means <- emmeans(model, run * laterality * subregion * affective_intensity * sensory_modality)

### Multivariate Pattern Analysis

We examined whether multivariate patterns within superior colliculus can distinguish the two sensory modalities. We compared multivariate patterns in superior colliculus between visual and somatosensory modalities using linear support vector machine (SVM) classifiers with a leave-one-subject-out cross-validation scheme (implemented in Nilearn; Abraham et al., 2014). SVM is a simple classification function that is widely used in neuroimaging, and its use eased the interpretation of models and minimized overfitting (Norman et al., 2006). First-level data from all runs for all subjects (except one subject; N = 79) in visual and somatosensory modalities were used to train classification models. Data from the one subject left out was used to test and estimate out-of-sample classification accuracy on their sensory modality label—i.e., could the left-out subject be correctly classified as completing a visual or somatosensory modality. The leave-out-subject-out cross-validation was repeated until each subject had been used for testing (80 folds in total). Classification accuracy was calculated based on the average accuracy score of each fold.

### Functional Connectivity Analysis

The analysis aimed to test whether signals in any brain regions correlated with signals in superior colliculus. The correlation among voxels was performed across trial-level estimates for all trials and runs within a subject (as opposed to model residuals within one run, as is standard in resting state connectivity analyses). This method was conducted using GLM estimated BOLD signal intensity from sensory stimulation in each trial, within each run, for each subject, consistent with both our pre-registered analysis plan (https://osf.io/pa5b9/) and prior work (see Kragel et al., 2021). Methods for regressor definition in visual and somatosensory modalities, nuisance parameters, and DARTEL warping to a common space were identical to those used in the group-level analysis, described above. No distinction in modeling was made between aversive and neutral trials—i.e., all trials were modeled as individual estimates. Trial-level whole-brain estimates were smoothed by using a 3 or 4 mm FWHM Gaussian kernel (for subcortical and cortical analysis/visualization, respectively). These trial-level whole-brain estimates were concatenated into a time series for each subject, providing a N-by-voxel map for each subject, where N = runs x trials per run (i.e., 5 x 24 if all runs were completed; in actuality, across subjects, for visual modality, M_N_ = 98.41, SD_N_ = 30.27; for somatosensory modality, M_N_ = 96.88, SD_N_ = 31.84). Connectivity between superior colliculus and the whole brain was estimated in each subject using a seed-based voxel-wise functional connectivity analysis. Superior colliculus signal was averaged for each trial within the subject-specific superior colliculus template, and trials were concatenated within each subject. Vectors for average superior colliculus signal and whole-brain signal within the grey matter mask were correlated to generate a whole-brain Pearson’s correlation coefficient map for each subject. Fisher’s r-to-z transformation was applied to this map to allow group-level analyses.

As stated in the pre-registration (https://osf.io/pa5b9/), we were concerned that stringent thresholding for the group analysis (e.g., voxel-wise *q*_FDR_ < .05) would increase false negatives (i.e., Type II error; Yarkoni, 2009), especially given that the superior colliculus is a small subcortical region with relatively noisy signals. To identify small, but reliable effects in both visual and somatosensory modalities, we developed a novel bootstrapping procedure, using group permutation to assess the reliability of connectivity. In this procedure, subjects were randomly divided into two separate sub-groups (N = 20 in each sub-group; see also, Kleckner et al., 2017; J. Zhang et al., 2019). In both groups, subject-level whole-brain connectivity maps (Fisher transformed) were submitted to a one-sample (OLS) t*-*test, estimating group-level whole-brain connectivity from SC. Voxels in both group maps were thresholded at *P* < .05, and the binarized conjunction of these two thresholded maps was saved. The procedure was performed separately for visual and somatosensory modalities, splitting the 40 subjects in each modality into two sub-groups of 20. For each modality, the procedure was repeated 1000 times, altering subject assignment into either sub-group in each iteration to produce 1000 binarized conjunction maps. For each modality, the 1000 group-level maps were converted to a percentage estimate, representing the reliability of a given voxel’s significant connectivity with superior colliculus. For example, a voxel estimate of 90% indicates that in 900 of 1000 conjunction maps, BOLD signal intensity in this voxel correlated with superior colliculus at *P* < .05 in two independent samples.

To calculate statistical significance for connectivity reliability, a null distribution was generated for each voxel. First, subject-level correlation coefficients (Fisher transformed) were independently shuffled, then, using the procedure described above, subjects for each modality were split into two equal groups, group-level whole-brain superior colliculus connectivity was estimated in each, and their binarized conjunction was saved. Initial subject-level voxel shuffling was performed 1000 times, and group assignments within each shuffle were performed 30 times, producing a null-distribution for each voxel comprised of 1000 observations. For each voxel, the observed reliability score was compared to the null distribution to compute a p value, and whole brain connectivity reliability was thresholded at -log_10_(*P*) > 1.3, corresponding to *P* < .05.

## Acknowledgements

This article was supported by grants from the National Cancer Institute (U01 CA193632 and R01CA258269-01), the National Science Foundation (BCS 1947972), the National Institute of Mental Health (R01 MH113234, R01 MH109464), the U.S. Army Research Institute for the Behavioral and Social Sciences (W911NF-16-1-019), the National Institutes of Health (NIA-R01AG063982), the Unlikely Collaborators Foundation, and the Roux Institute and the Harold Alfond Foundation. The views, opinions, and/or findings contained in this review are those of the author and shall not be construed as an official Department of the Army position, policy, or decision, unless so designated by other documents, nor do they necessarily reflect the views of the Unlikely Collaborator Foundation.

## Supplementary Materials

**Fig S1.**
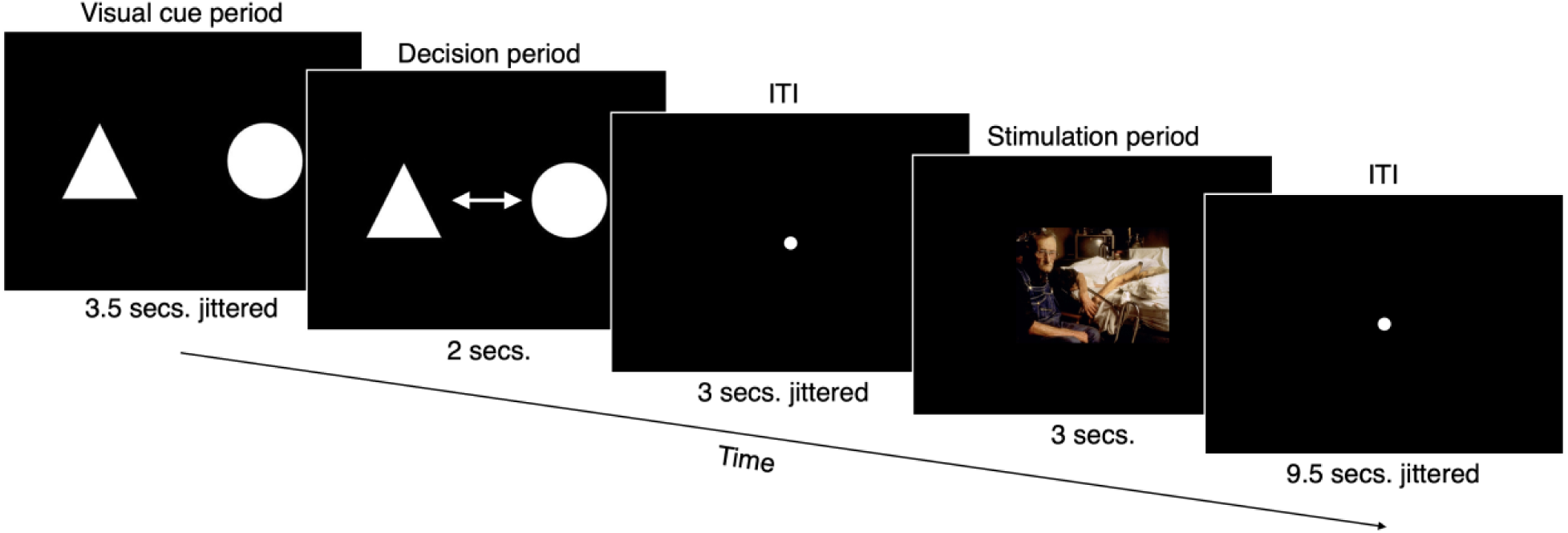
Experiment paradigm for one trial. The experiment was originally designed as a probabilistic avoidance learning task. In the visual cue period participants viewed two visual cues (sampled from Poisson distribution, 3.5 secs. on average). In the decision period (2 secs.), participants were either asked to choose one of the shapes (50% of trials) or a shape was randomly chosen for them (the remaining 50% of trials). After a jittered inter-trial interval (ITI; 1-5 secs., 3 secs. on average), either an image was displayed (in visual stimulation; 3 secs.), or mechanical pressure was delivered to the left thumb (in somatosensory stimulation; 3 secs). Images were from the IAPS database (Lang et al., 1997) and normatively rated to be either negative or neutral valence, while mechanical pressure was calibrated at the beginning of the scan to be either high or low pressure. Aversive or neutral stimulation in either stimulation was probabilistically related to the choice of shape made in the decision period, with probabilities changing in a random walk across the experiment (see Methods). Note that, during the somatosensory stimulation, a fixation point was displayed (in place of an image) during stimulation period. Trials were separated by a jittered ITI (sampled from Poisson distribution, 9.5 secs. on average). Only the stimulation period was relevant to the present work.

**Fig S2.**
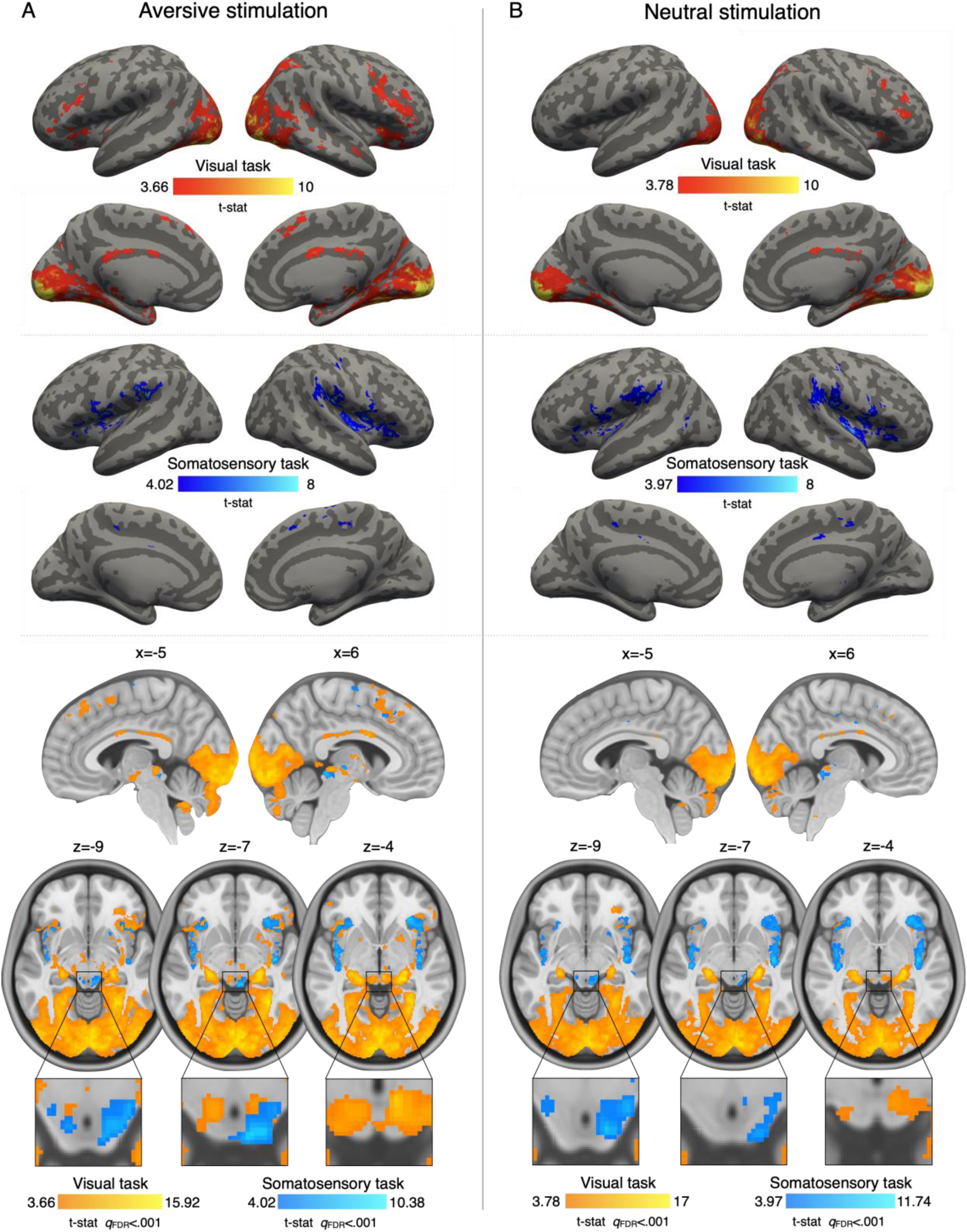
Group-level whole brain univariate analysis for (A) aversive stimulation, and (B) neutral stimulation, during visual (orange) and somatosensory (blue; on top of orange in the lowest panel) stimulation, relative to baseline. Both panels report one-tailed *t* statistics (stimulation > baseline), thresholded at a voxel-wise cutoff of *q*_FDR_ < 0.001 and cluster size k >= 20 voxels. During visual stimulation, aversive stimulation (negative valence-normed images) elicited BOLD intensity in dorsal anterior insula, temporal fusiform gyrus, paracingulate gyrus, mid to posterior cingulate cortex, orbital frontal cortex, occipital cortex, amygdala, right hippocampus, mammillary nuclei, putamen, caudate, posterior pulvinar, lateral geniculate nuclei, subthalamic nucleus, red nucleus, and the substantia nigra (in addition to dorsal superior colliculus, see Fig. 1). During somatosensory stimulation, aversive stimulation (high mechanical pressure) elicited BOLD activity in, as well as in the anterior to posterior insula, right ventroposterior thalamic nuclei, paracingulate gyrus, midcingulate cortex, and right primary somatosensory cortex (in addition to ventral superior colliculus, see Fig. 1). Neutral valence-normed images and low mechanical pressure elicited activity in similar brain regions, but generally to a lesser extent.

### Multivariate patterns in superior colliculus distinguished between visual and somatosensory stimulation

Multivariate pattern analysis provided an additional, more sensitive test of visual–somatosensory differences in superior colliculus activity. The results indicated that visual and somatosensory stimulation could be distinguished by voxel-wise patterns in the superior colliculus. A linear SVM classifier with leave-one-subject-out cross-validation was trained on voxel-wise patterns of superior colliculus BOLD intensity using both aversive and neutral run-level estimates. The trained classifiers distinguished visual and somatosensory stimulation labels significantly above chance level, *t*_(79)_ = 12.06, *P* < .001, with mean accuracy of 79.42% and standard error of 2.44%. The classifiers continued to perform above chance when limited to only aversive (*t_(79)_* = 7.64, *P* < .001; mean accuracy = 75.31%; standard error = 3.31%) or neutral (*t_(79)_* = 7.64, *P* < .001; mean accuracy = 75.31%; standard error = 3.31%) run-level estimates. The identical accuracies and standard errors in both aversiveness conditions were a coincidence.

**Table S1.** Linear mixed effects model summary. F-values and P-values in the table used Satterthwaite’s method for denominator degrees-of-freedom and F-statistic. For the details of model description, see Methods. *** denotes *P* < .001; ** denotes *P* < .01; * denotes *P* < .05.

|  | Sum of squares | Mean square | Numerator df | Denominator df | F value | *P* |
| --- | --- | --- | --- | --- | --- | --- |
| run | 1794.5 | 448.6 | 4 | 64.152 | 1.6768 | 0.16623 |
| laterality | 6158.0 | 6158.0 | 1 | 159.676 | 23.0174 | 3.659e-06 *** |
| subregion | 11952.1 | 11952.1 | 1 | 79.513 | 44.6747 | 2.917e-09 *** |
| affective_intensity | 1443.9 | 1443.9 | 1 | 66.789 | 5.3972 | 0.02322 * |
| sensory_modality | 1670.8 | 1670.8 | 1 | 70.160 | 6.2451 | 0.01480 * |
| laterality: affective_intensity | 272.4 | 272.4 | 1 | 257.823 | 1.0183 | 0.31387 |
| subregion affective_intensity | 18.1 | 18.1 | 1 | 176.932 | 0.0676 | 0.79513 |
| run: sensory_modality | 1282.5 | 320.6 | 4 | 64.152 | 1.1984 | 0.32014 |
| laterality: sensory_modality | 4860.3 | 4860.3 | 1 | 159.676 | 18.1669 | 3.45e-05 *** |
| subregion: sensory_modality | 22571.6 | 22571.6 | 1 | 79.513 | 84.3683 | 4.004e-14 *** |
| affective_intensity: sensory_modality | 47.7 | 47.7 | 1 | 66.789 | 0.1784 | 0.67411 |
| laterality: affective_intensity: sensory_modality | 465.8 | 465.8 | 1 | 257.823 | 1.7412 | 0.18815 |
| subregion: affective_intensity: sensory_modality | 75.4 | 75.4 | 1 | 176.932 | 0.2819 | 0.59612 |

**Fig S3.**
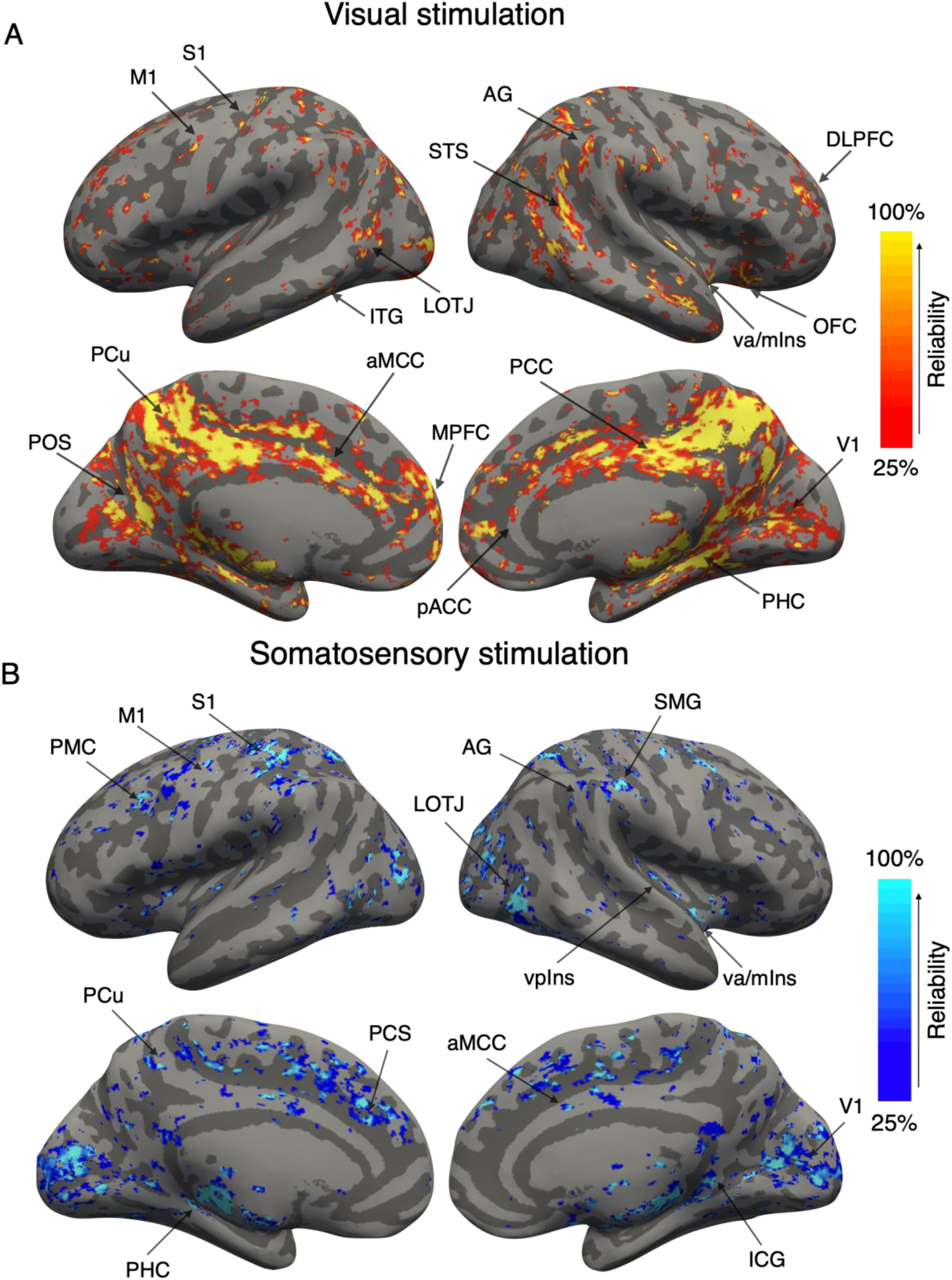
Group-level whole-brain reliability of functional correlations with superior colliculus activity during (A) visual (orange) and (B) somatosensory (blue) stimulation. Reliability was estimated the percentage of overlap in a bootstrapped split-half procedure (1000 iterations; see Methods). Both panels show reliability thresholded at 25% and cluster size of k >= 20 voxels. Data was smoothed by a 4mm FWHM Gaussian kern. Unthresholded maps are available online: https://neurovault.org/collections/MKTVBWGR/. Abbreviations: S1: primary somatosensory cortex; M1: primary motor cortex; AG: angular gyrus; STS: superior temporal sulcus; vaIns: ventral anterior insula; OFC: orbital frontal cortex; PCu: precueus cortex; aMCC: anterior mid-cingulate cortex; MPFC: medial prefrontal cortex; pACC: pregenual anterior cingulate cortex; PCC: posterior cingulate cortex; V1: primary visual cortex; PHC: parahippocampal cortex; PCS: paracingulate sulcus; SMG: supramarginal gyrus; LOTJ: lateral occipito-temporal junction; ICG: isthmus of cingulate gyrus.

**Fig S4.**
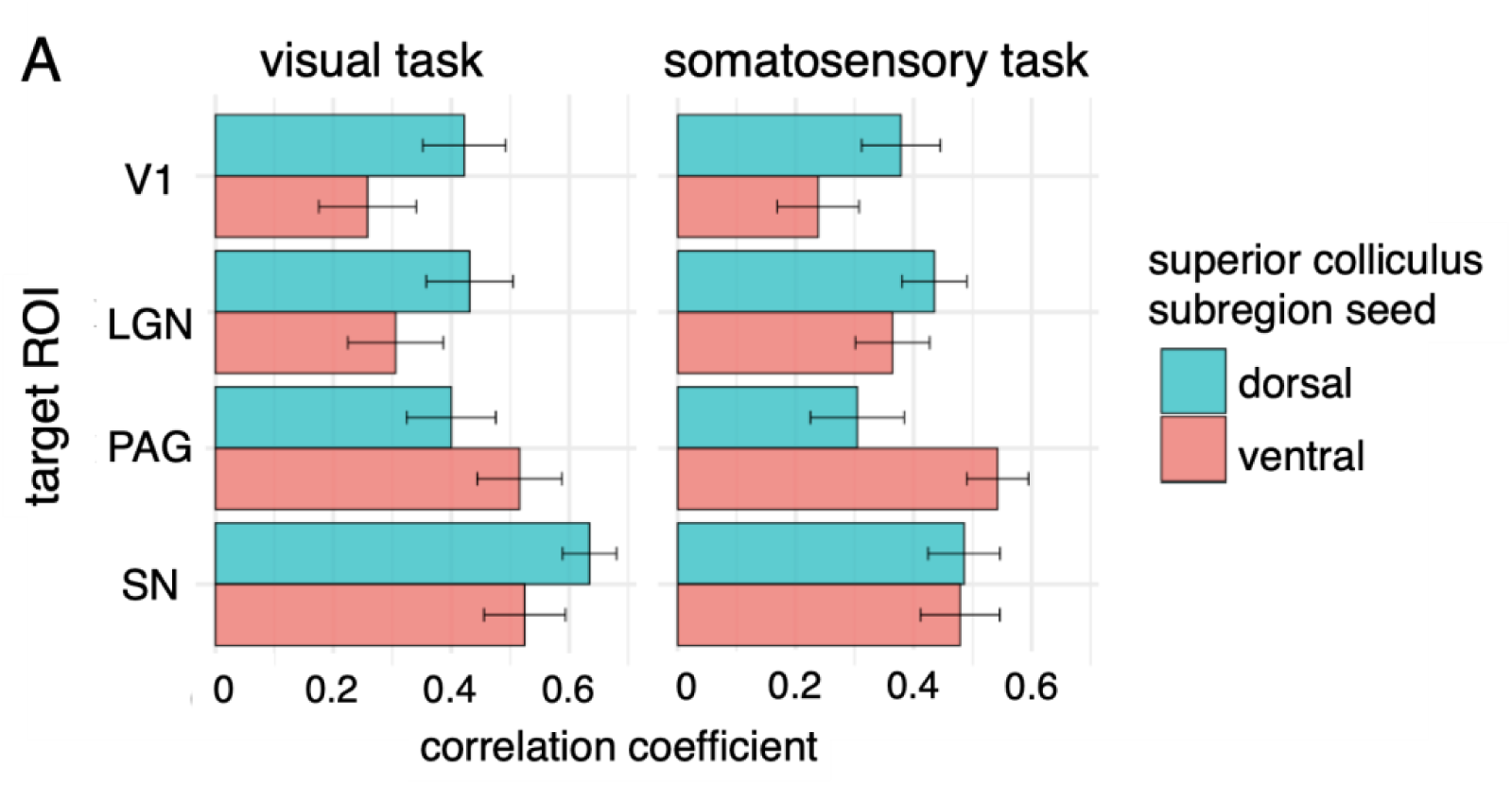
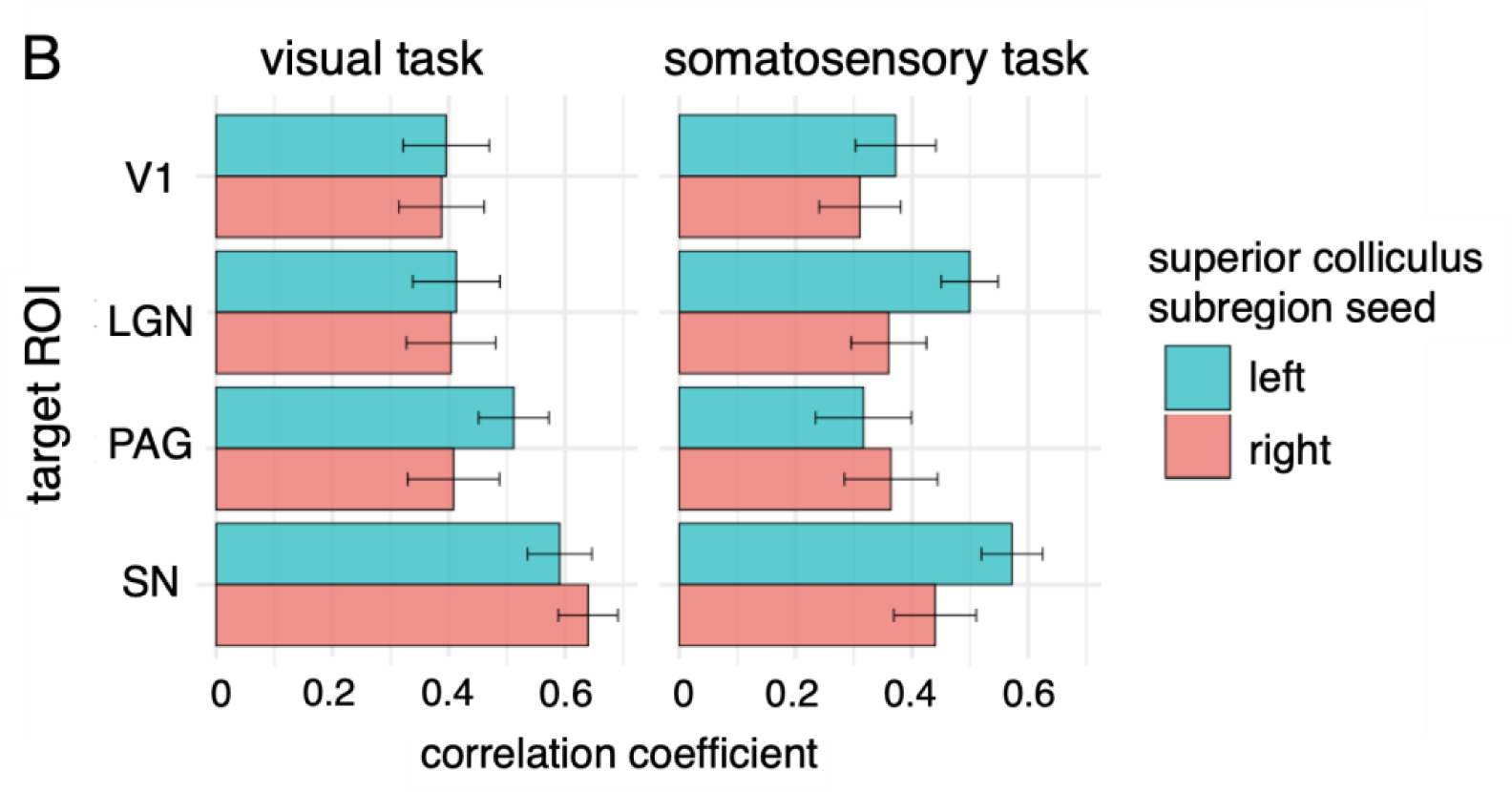
ROI-to-ROI functional connectivity between superior colliculus subregion seeds and selected targets (lateral geniculate nucleus (LGN), periaqueductal grey (PAG), substantia nigra (SN), and primary visual cortex (V1)). The analysis correlated mean signals of superior colliculus subregions with mean signals of entire target regions from each subject’s concatenated trial-level estimates (same as data used in other functional connectivity analyses; see Methods). (A) Dorsal superior colliculus signals showed greater correlation with traditionally considered visual regions such as V1 and LGN, compared to ventral superior colliculus signals, for both sensory modality stimulation, consistent with anatomical connectivity evidence. Similarly, ventral superior colliculus signals showed greater correlation with PAG, compared to dorsal superior colliculus signals, for both sensory modality stimulation, consistent with anatomical connectivity evidence. Dorsal superior colliculus signals showed greater correlation with SN during visual stimulation, while dorsal and ventral superior colliculus did not differ in their correlation with SN during somatosensory stimulation. (B) Left and right superior colliculus did not show significant difference with target regions during visual stimulation. Left superior colliculus showed greater correlation with LGN and SN. Future work is needed to explore this difference.

## Footnotes

1 Functional connectivity analyses in the present study used the BOLD signal intensity from the entire superior colliculus, as greater number of voxels within the superior colliculus, a small region, is expected to yield more reliable results. For superior colliculus subregion connectivity with select targets, see Fig. S4.

2 Due to power concerns, we did not perform separate functional connectivity analyses for aversive and neutral affective intensity conditions in each sensory modality condition (for more on the importance of within-subject sample sizes, see Gonzalez-Castillo et al., 2012; for suggested length of sliding window in dynamic functional connectivity approach, see Savva et al., 2019).

